## Supplementary Information for "Deep-Learning Structure Elucidation from Single-Mutant Deep Mutational Scanning"

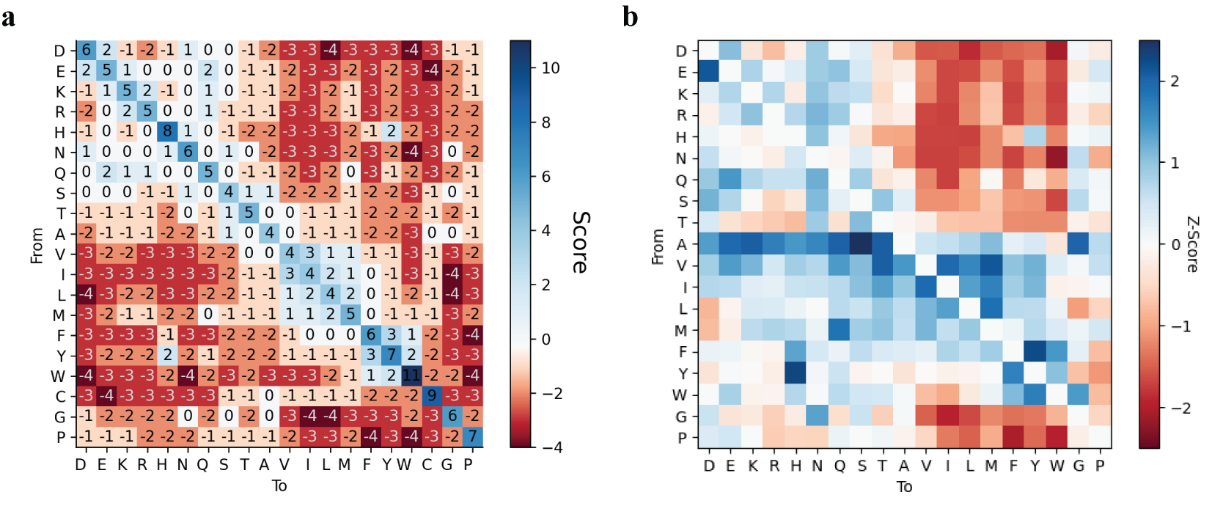

**Supplementary Figure 1)** a) BLOSUM62 scoring matrix. High and low BLOSUM scores indicate whether a mutation is likely conservative or nonconservative, respectively. b) Z-scores of all residue substitution types between normalized BLOSUM62 scores and mutational correlations derived from comparing burial extent and ΔΔGs of the mega-scale dataset. A high Z-score means a particular mutation is more likely to be detrimental according to our analysis. It corresponds to a higher BLOSUM score but exhibits a stronger mutation correlation. A low Z-score means a particular mutation is more likely to be detrimental according to BLOSUM. It corresponds to a lower BLOSUM score but has a weaker mutation correlation.

| Wild Type Residue | Mutated Residue | $\boldsymbol{R}^{\boldsymbol{2}}$ | Wild Type Residue | Mutated Residue | $\boldsymbol{R}^{\boldsymbol{2}}$ |
| --- | --- | --- | --- | --- | --- |
| A | N | 0.48 | **V** | N | 0.41 |
| A | K | 0.47 | **I** | D | 0.41 |
| A | Q | 0.47 | **A** | R | 0.41 |
| A | E | 0.45 | **I** | Q | 0.40 |
| A | H | 0.45 | **I** | E | 0.40 |
| V | Q | 0.44 | **V** | K | 0.40 |
| W | G | 0.44 | **V** | T | 0.39 |
| V | E | 0.44 | **V** | S | 0.39 |
| A | D | 0.44 | **I** | N | 0.38 |
| V | H | 0.43 | **A** | T | 0.38 |
| V | D | 0.43 | **I** | S | 0.38 |
| A | S | 0.43 | **I** | H | 0.38 |
| W | E | 0.42 |  |  |  |

**Supplementary Table 1)** Top 25 mutational types with the highest correlation between mutational stability and burial extent.

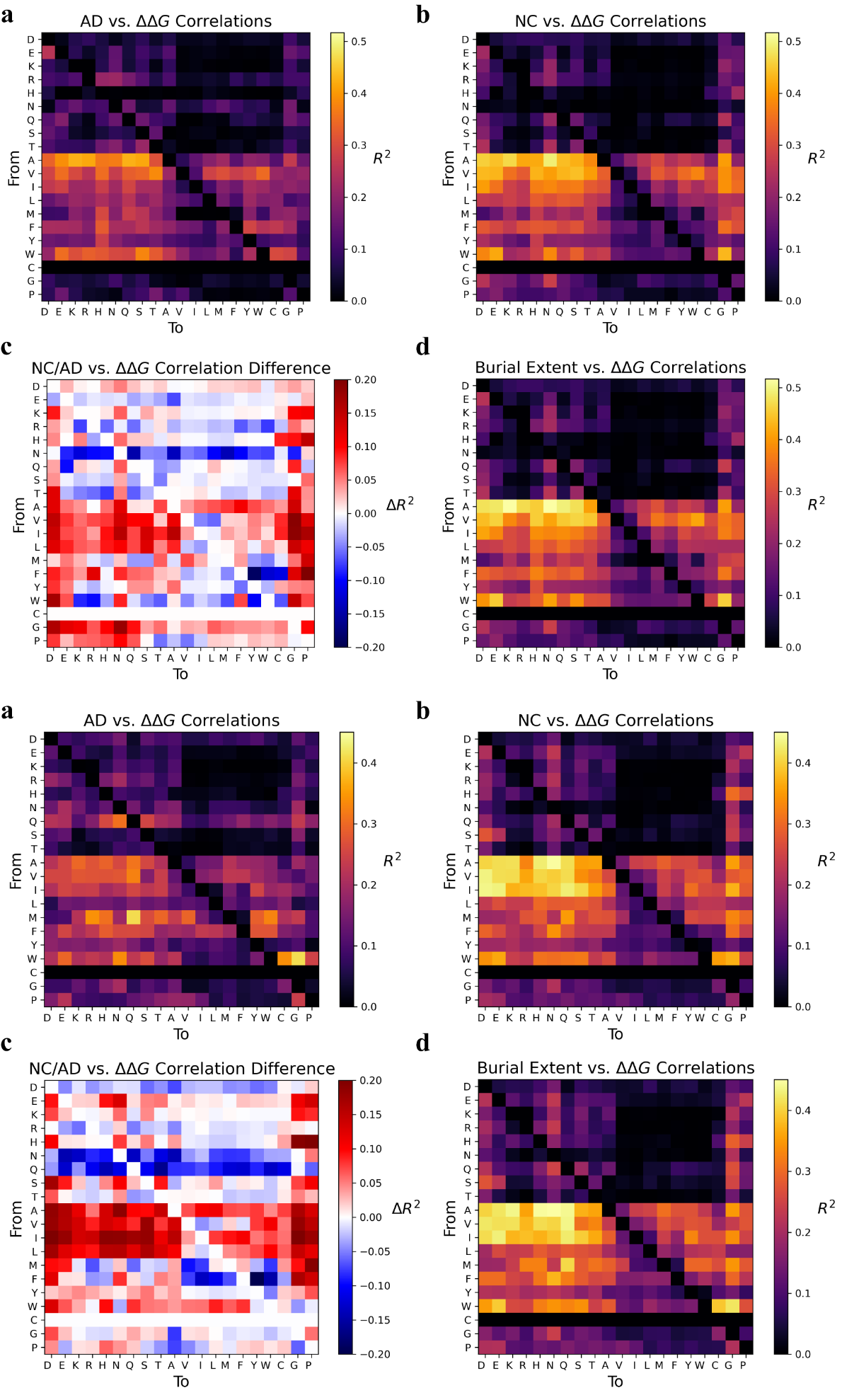

**Supplementary Figure 2)** Heat maps depicting the correlation between ΔΔG and solubility metrics for individual mutational types across the first mega-scale subset (88 proteins). a) Comparing ΔΔGs to native residue atomic depth. b) Comparing ΔΔGs to native neighbor count. c) Differences in correlation coefficients between atomic depth and neighbor count. d) Comparing ΔΔGs to burial extent (defined as weighted average of both atomic depth and neighbor count).

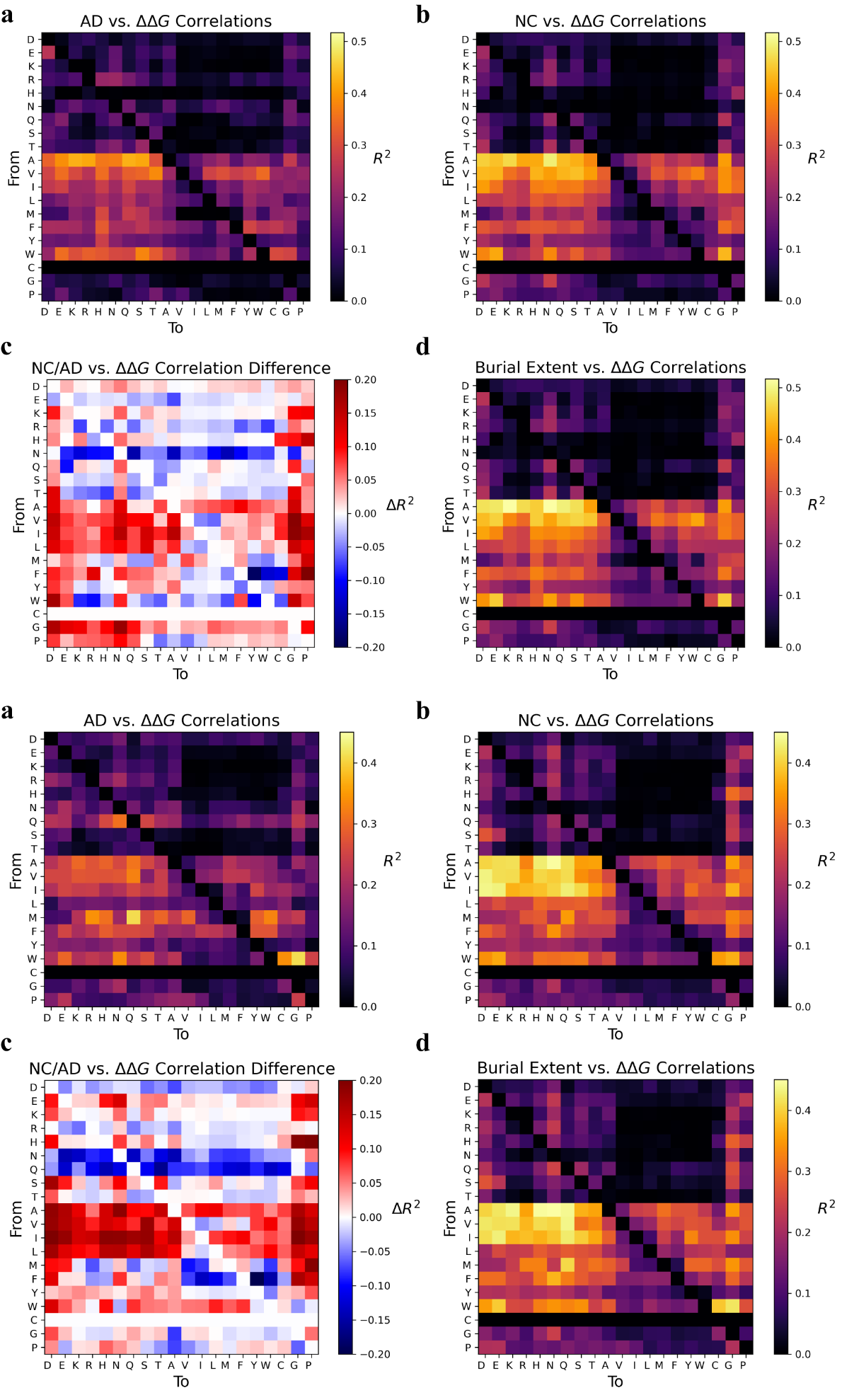

**Supplementary Figure 3)** Heat maps depicting the correlation between ΔΔG and solubility metrics for individual mutational types across the second mega-scale subset (87 proteins). a) Comparing ΔΔGs to native residue atomic depth. b) Comparing ΔΔGs to native neighbor count. c) Differences in correlation coefficients between atomic depth and neighbor count. d) Comparing ΔΔGs to burial extent (defined as weighted average of both atomic depth and neighbor count).

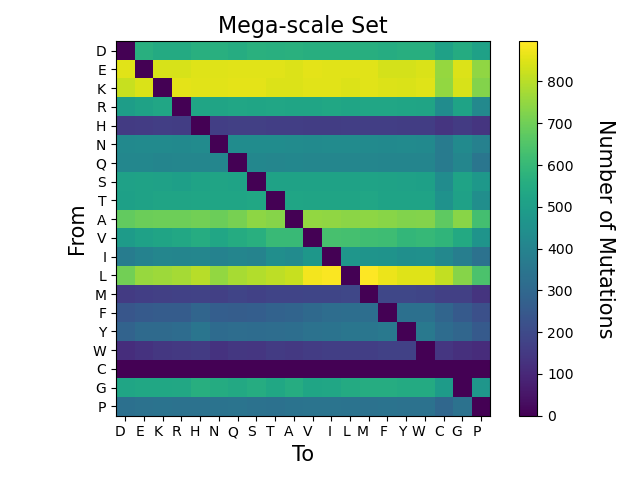

**Supplementary Figure 4)** Number of mutations across the 175 proteins taken from the mega-scale set. Due to the low number of cysteine residue mutations, these were excluded from our analysis.

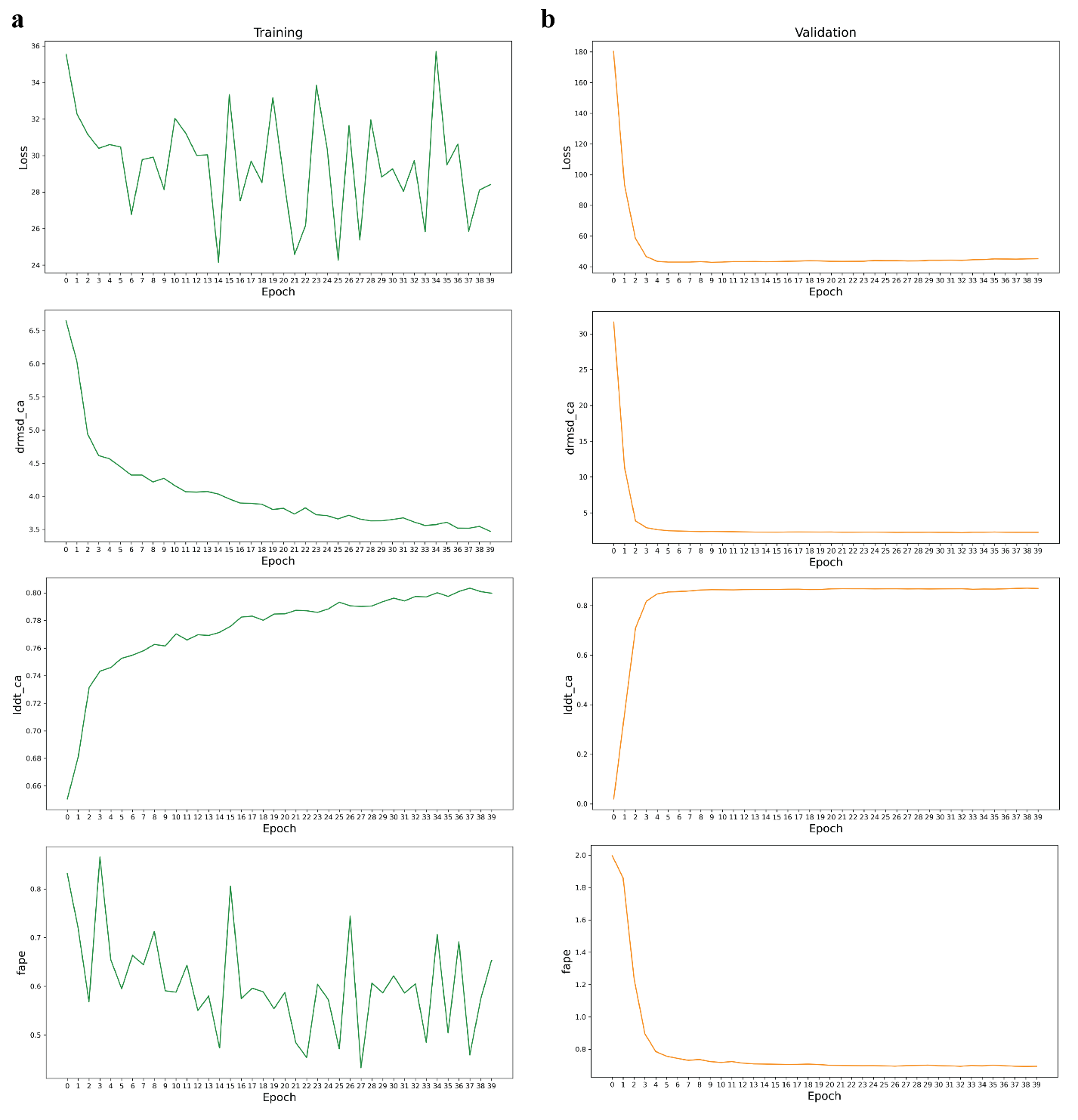

**Supplementary Figure 5)** a) Subset of DMS-Fold training metrics, including training loss per-epoch, drmsd_ca per-epoch, lddt_ca per-epoch, and fape loss per-epoch. b) Subset of DMS-Fold validation metrics, including validation loss per-epoch, drmsd_ca per-epoch, lddt_ca per-epoch, and fape loss per-epoch.

**Supplementary Table 2)** PDB IDs of proteins in each subset of the mega-scale set.

Set 1 (88 Proteins)

1GYZ, 1UZC, 2MH8, 1I6C, 6ACV, 2M8E, 5UYO, 2K5P, 1W4G, 2LVN, 2JWT, 2M8J, 2L2P, 2MC5, 2GP8, 2L2D, 2LQK, 2WNM, 2KXD, 6FVC, 6OBK, 2MCK, 1GL5, 1O6X, 1JIC, 5UP5, 2O2W, 2B89, 2JWS, 2KZJ, 1W4H, 2M2J, 1UFM, 2K5N, 3I35, 2LYQ, 2LHR, 5Z2S, 2LC2, 2BTT, 4G3O, 2L09, 3CQT, 7JJK, 1QLY, 2M8U, 2K2A, 2OCH, 6YSE, 7BPM, 3ZGK, 1LP1, 1TUC, 2MA4, 2KRU, 2N4R, 3DKM, 1VII, 2RU9, 1W4F, 2K5H, 2N88, 2JVG, 1GJS, 6EWU, 2JVD, 1PV0, 5OAO, 1IFY, 2KVS, 1YP5, 2KFV, 3MYC, 2D1U, 5LXJ, 2YSB, 1YRF, 2WQG, 1PSE, 1QKX, 2LUM, 2KCM, 1QP2, 2KGT, 1R69, 2K9D, 2L7F, 2JN4

Set 2 (87 Proteins)

1K1V, 2LHC, 6EWT, 1V1C, 2M9F, 2M0C, 1ZLM, 2AMI, 2MYX, 3L1X, 1TUD, 2JT1, 1WCL, 1YU5, 2OP7, 2KVT, 1F0M, 1I2T, 2M8I, 6M3N, 6SCW, 2B88, 5JRT, 2L33, 4C26, 1URF, 6IWS, 2BTH, 2M6Y, 2MI6, 3V1A, 1WR4, 2L9R, 1QKH, 5VNT, 2K1B, 4UZW, 2KCF, 1Y0M, 2YSF, 2EXD, 5UBS, 1TG0, 2CJJ, 6NMW, 2M9I, 2M2L, 1A32, 6SOW, 1PWT, 2HBB, 2L7M, 1H92, 2M0Y, 2L6Q, 1ORC, 2JTV, 2M9E, 5UCE, 2RJV, 1SRM, 1E0L, 2LYR, 2RRU, 1ZHC, 6EWS, 2LCL, 2LGW, 2QFF, 2KR3, 2RRT, 2MKY, 2MXD, 2K28, 5AHT, 2LO1, 1ENH, 2LP5, 2MCH, 1OPS, 2WXC, 1AOY, 2LJ3, 2JZ2, 4HCK, 5KPH, 2KWH
